# A proximity labeling approach to identify proteins that associate with synaptonemal complex components in *Drosophila melanogaster* females

**DOI:** 10.1101/2025.10.14.682398

**Authors:** Stacie E. Hughes, Christopher Viermann, Morgan James, Charles Banks, R. Scott Hawley

## Abstract

Organisms use a specialized cell division called meiosis for the creation of haploid gametes. Multiple carefully orchestrated steps must occur at specific times and places for meiosis to be successful, including chromosome pairing, meiotic entry, recombination, synapsis, and two rounds of chromosome segregation. The regulation and molecular mechanisms for many of the steps of meiosis have not been fully elucidated. During synapsis, the synaptonemal complex (SC) builds along the entire lengths of the homologs to maintain the pairing of the homologs and promote the formation of the crossovers that help ensure proper segregation of homologs at the meiosis I division in many organisms. The SC is a large tripartite structure that is believed to function as a biomolecular condensate. To attempt to identify proteins that interact with SC components during female meiosis in *Drosophila melanogaster*, a protein of the lateral element, C(2)M, and a protein of the central element, Cona, were tagged with the APEX2 enzyme, which can biotinylate nearby proteins under the appropriate conditions. Under biotinylating promoting conditions, biotin labeled proteins were observed to be associated with the SC by immunofluorescence. Biotinylated proteins were isolated for mass spectrometry analysis, and multiple proteins were found to be enriched compared to control samples. RNAi knockdown lines targeting a subset of enriched proteins were examined for phenotypes in early *Drosophila* female meiosis. RNAi knockdown of *Cpsf5*, an mRNA cleavage factor, caused delayed or defective SC formation, as well as additional meiotic defects, indicating a role for maturation of mRNA in regulating processes of female meiosis. These results support proximity labeling as a strategy for identifying additional meiotic proteins.

## Introduction

Sexual reproduction requires the production of haploid gametes that are made through meiosis, a specialized cell division program. For meiosis to be successful, multiple events must occur at specific times and places in the germline and these events must be coordinated with gamete development. In a canonical meiosis, cells enter the meiotic program, pair chromosomes, induce numerous double strand breaks (DSBs), form the synaptonemal complex (SC), repair the DSBs as either crossovers or noncrossovers, segregate homologs away from each other at the first division, and finally segregate sister chromatids at the second division. Many organisms rely on the formation of crossovers between the homologous chromosomes to ensure homologs segregate away from each other at the first division. Defects in early meiotic events can lead to defective gametes or subsequent failures in chromosome segregation that ultimately lead to aneuploid offspring or lethality. Despite the importance of this process, many mechanistic aspects of meiosis are still poorly understood.

The SC was identified by electron microscopy decades ago (Moses 1968) and the composition, structure, and function have been of great interest. Besides maintaining the association of homologs, the SC is required for crossing over in many organisms, and recent evidence suggests the SC influences crossover patterning (Capilla-Perez *et al*. 2021; Hughes *et al*. 2025; Libuda *et al*. 2013; Williams *et al*. 2025). The SC has been proposed to behave as a biomolecular condensate that influences crossover placement through the coarsening of pro-crossover factors at a subset of DSBs (Durand *et al*. 2022; Girard *et al*. 2023; Von Diezmann *et al*. 2024). By electron microscopy, the SC has a ladder-like structure with two lateral elements (LE) associated with the homologous chromosomes and a central region that connects the LEs. In *Drosophila* females, the LEs are composed of cohesin and cohesin-related proteins, including the C(2)M protein (Fig 1). C(2)M has been proposed to work with Nipped-B and stromalin and has been shown to physically interact with SMC3 in two-hybrid studies (Fig. 1) (Anderson *et al*. 2005; Gyuricza *et al*. 2016; Heidmann *et al*. 2004). Loss of C(2)M leads to a failure of the SC to extend into full-length tracks after the initial loading of SC central region components at centromeres and a small number of additional sites along the chromosomes, as well as, defects in the levels of crossing over (Manheim and Mckim 2003). The known proteins of the central region are the transverse filament protein C(3)G, Corolla, and Cona (Collins *et al*. 2014; Page and Hawley 2001; Page *et al*. 2008) (Fig 1). All three proteins are necessary for full-length SC formation, and null mutations eliminate crossing over, cause defective centromere clustering, and lead to high levels of chromosome missegregation (Collins *et al*. 2014; Page and Hawley 2001; Page *et al*. 2008; Takeo *et al*. 2011).

**Figure 1.**
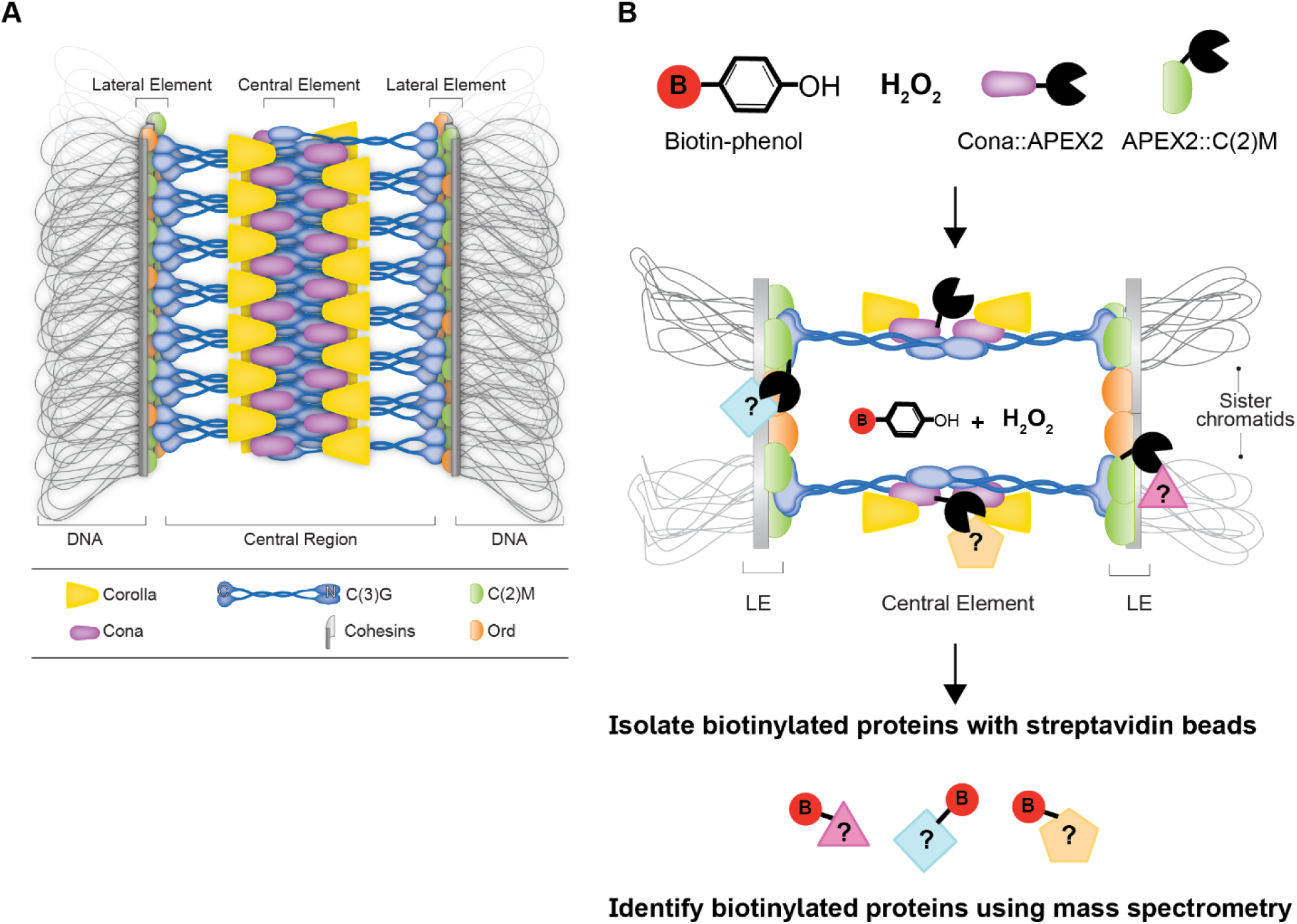
Model of the *Drosophila* SC and schematic of the proximity labeling experiment. (A) Shown is the current model of the *D. melanogaster* SC. (B) Top view model of the *D. melangaster* SC with a schematic of the proximity ligation by APEX2::C(2)M and Cona::APEX2. APEX2::C(2)M and Cona::APEX2 incorporate into the LE and central region of the SC. In the presence of biotin-phenol and H2O2 APEX2 transfers biotin to electron-rich amino acids on nearby proteins. Nuclei are isolated from biotin-phenol treated ovaries and the biotinylated proteins were isolated on streptavidin coated beads for mass spectrometry analysis. Adapted from (Cahoon *et al*. 2017).

Identifying the proteins involved in meiotic processes can be challenging. Many meiotic genes, including SC components, are rapidly evolving and cannot be identified using only simple homology searches to more distant organisms (Hemmer and Blumenstiel 2016; Kursel *et al*. 2021; Williams and Hawley 2025; Zakerzade *et al*. 2025). Forward genetic screens were the early method used to discover new genes involved in meiosis in *D. melanogaster* females (reviewed in (Hughes *et al*. 2018)). Besides being labor intensive, the screens relied on mutagens that induce only specific types of mutations, and the screens mostly selected for strong meiotic phenotypes for the identification of the new mutations. As these screens relied on the viability and fertility of females to be recovered, genes with additional roles in adult viability or mutations that strongly impair meiosis or oogenesis would fail to be recovered.

To try to identify those classes of genes which would have been missed in forward genetic screens, efforts have turned to RNAi based screens, which eliminate potential issues with adult viability and some types of sterility. These screens have either been mostly unbiased in the selection of the lines tested (Gilliland *et al*. 2024; Nieken *et al*. 2023) or candidates chosen based on gene expression (Hughes *et al*. 2024; Sun *et al*. 2024). These efforts provided important information on additional roles of known genes in meiotic regulation and have yielded new candidates for future studies.

Proteins that localize near known meiotic proteins, such as SC proteins, would be strong candidates for also playing roles in meiosis. Recent studies have used proximity labeling approaches to identify novel proteins of biological processes and to gain additional information on protein interactions. One method of proximity labeling is to use a form of ascorbate peroxidase enzyme (APEX) to biotinylate proteins in close physical proximity to tyrosine and other electron-rich amino acids in the presence of biotin-phenol and hydrogen peroxide (Hung *et al*. 2016; Kim and Roux 2016). APEX enzymes have high enzymatic activity at a wide range of temperatures under relatively short labeling reaction times (seconds to minutes). One technical challenge of using APEX enzymes is the requirement for biotin-phenol, which has low membrane permeability. APEX fused proteins were successfully utilized in the *D. melanogaster* ovary to study interactions among ring canals proteins within egg chambers by using detergent to improve biotin-phenol penetration (Mannix *et al*. 2019). A mutated version of the APEX enzyme was identified, called APEX2, which contains a mutation (A134P) that improves enzyme kinetics and thermal stability (Lam *et al*. 2015). By tagging two proteins of the *Drosophila* SC with the APEX2 enzyme we hoped to identify potentially new SC proteins that may play roles in key steps of meiosis.

We tagged a LE (C(2)M) and a central region (Cona) component of the SC with the APEX2 enzyme. Ovaries expressing the *c(2)M* (*APEX2::c(2)M*) or *cona* (*cona::APEX2*) APEX2 tagged constructs with a germline specific *nanosGAL4::VP16* driver exposed to biotin-phenol labeling conditions displayed biotin labeling near the SC in an immunofluorescence assay. Biotinylated proteins from the *APEX2::c(2)M* and *cona::APEX2* expressing ovaries were isolated, along with proteins from biotin-phenol treated controls, and sent for mass spectrometry analysis. While some proteins with known roles in the *D. melanogaster* female ovary were found enriched in the *APEX2::c(2)M* or *cona::APEX2* samples, additional proteins were found that had only limited information, if any, on their role in the female germline or female meiosis. As an initial investigation into some of the lesser characterized enriched proteins, germline expressed shRNAi lines generated by the TRiP consortium were analyzed by immunofluorescence for defects in SC and DSB dynamics. We identified that germline expression of a shRNA targeting *CG3689* (*Cleavage and polyadenylation specific factor 5* (*Cpsfs5*)) caused defects in full-length SC formation, chromosome segregation, and crossover placement. Cpsf5 is predicted to be part of an mRNA cleavage factor complex. Our results suggest Cpsf5 may regulate aspects of female meiotic progression. Using the strategy of investigating proteins in proximity to known meiotic proteins has the potential to yield new information on the regulation and mechanism of meiotic processes.

## Results and Discussion

To identify new meiotic proteins, we used an approach that examined proteins associated with or in close physical proximity to two proteins of the SC. The regulation of the assembly and disassembly of the SC in *Drosophila* females is still poorly understood, especially because the *Drosophila* SC has not been reconstituted in vitro. The LE protein C(2)M and the central region protein Cona were chosen for fusion to APEX2 since they are believed to occupy two different parts of the SC (Fig. 1) (Anderson *et al*. 2005; Cahoon *et al*. 2017), and tagged versions of the proteins had been successfully used in previous studies (Manheim and Mckim 2003; Page *et al*. 2008). Two inducible germline expression constructs were created by fusing APEX2 to the N-terminus of C(2)M and the C-terminal end of Cona in a *pUASp* expression vector creating *APEX2::c(2)M* or *cona::APEX2*, respectively. Tagging location was based on the success of previous tagged versions of the proteins (Manheim and Mckim 2003; Page *et al*. 2008). We opted to use an inducible germline vector allowing for expression only in the generation of interest to limit possible issues in stock maintenance due to the large tags inhibiting the function of the SC proteins. The constructs were driven by the *nanos-GAL4::VP16* driver, which drives robust expression of *pUASp* constructs in the female germline (Rorth 1998). Expression of the *APEX2::c(2)M* or *cona::APEX2* with *nanos-GAL4::VP16* in the background of untagged copies of the SC proteins did not appear to cause any dominant detrimental meiotic phenotypes in regard to SC assembly in pachytene based on localization of the transverse filament protein C(3)G or on chromosome segregation (Table 1, Fig 2). As a control for non-specific biotin labeling by APEX2 tagged proteins during protein production and transport, as well as from the overexpression conditions, an APEX2 only construct was created in *pUASp* (*APEX2*) for comparisons in proteomics analysis.

**Figure 2.**
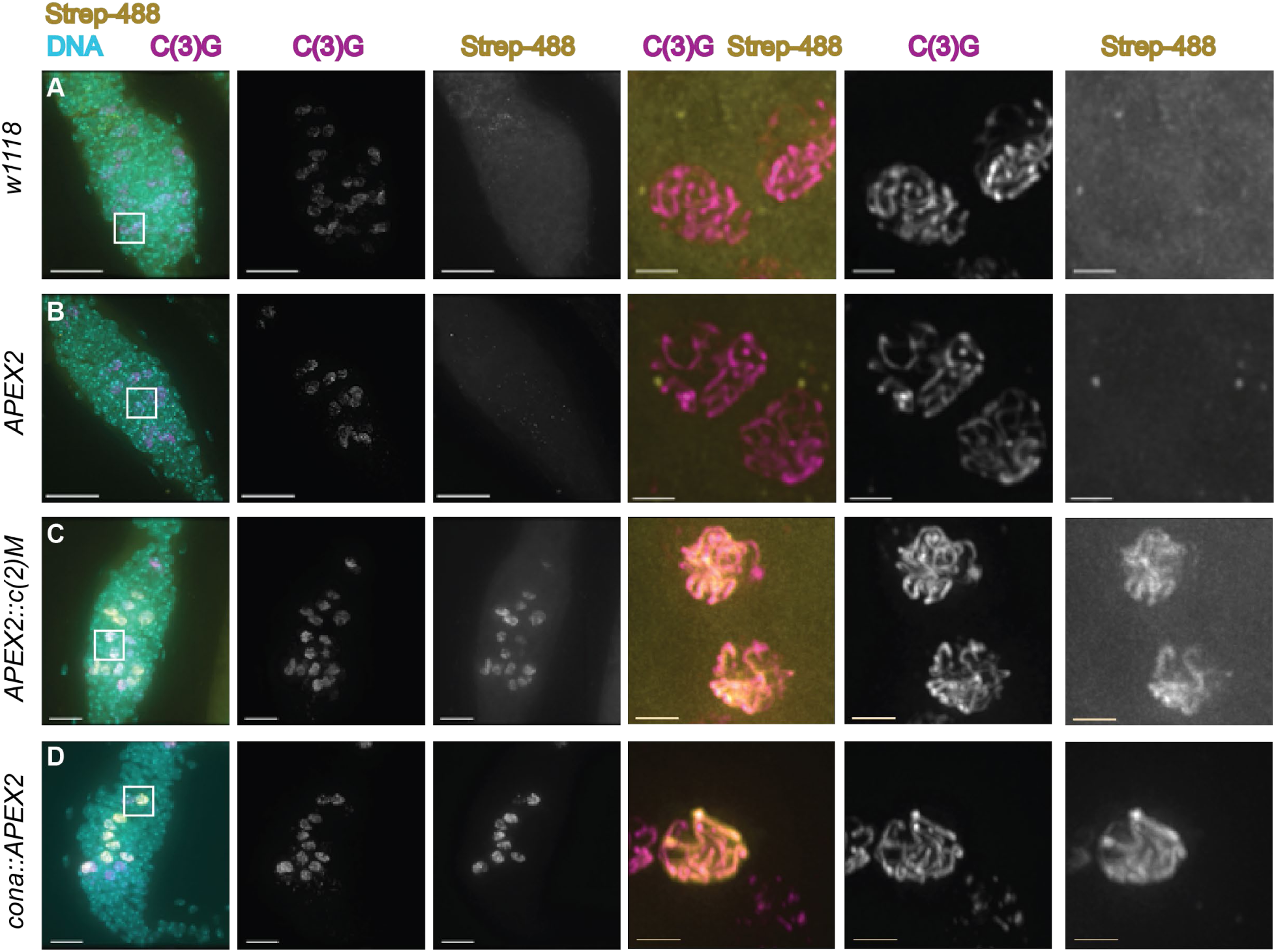
Proteins in close proximity to the SC are biotinylated in *cona::APEX2* and *APEX2::c(2)M* expressing flies. Ovaries underwent biotin-phenol labeling followed by immunofluorescence with an antibody against C(3)G (SC, magenta), strevadin-488 (recognizes biotin, yellow), and DAPI (DNA, cyan). Germaria from (A) *w^1118^* and females expressing *APEX2* did not show similar localization patterns between streptavidin-488 and C(3)G. In *APEX2::c(2)M* (C) and *cona::APEX2* (D) germaria streptavidin localized in a pattern similar to full-length SC in a subset of nuclei. Boxed regions in the first column are shown enlarged without DAPI. Images are projections from z-stacks. Scale= 10 µm in germaria images and 2 µm in enlarged nuclei images. Full genotypes are *w^1118^*, *y w; pUASp-APEX2/+; nanosGAL4::VP16/+; spa^pol^/+*, *y w;; pUASp-APEX2::c(2)M; nanosGAL4::VP16/+; spa^pol^/+*, and *y w; pUASp-cona::APEX2/+; nanosGAL4::VP16/cona^f04903^; spa^pol^/+*.

**Table 1.**
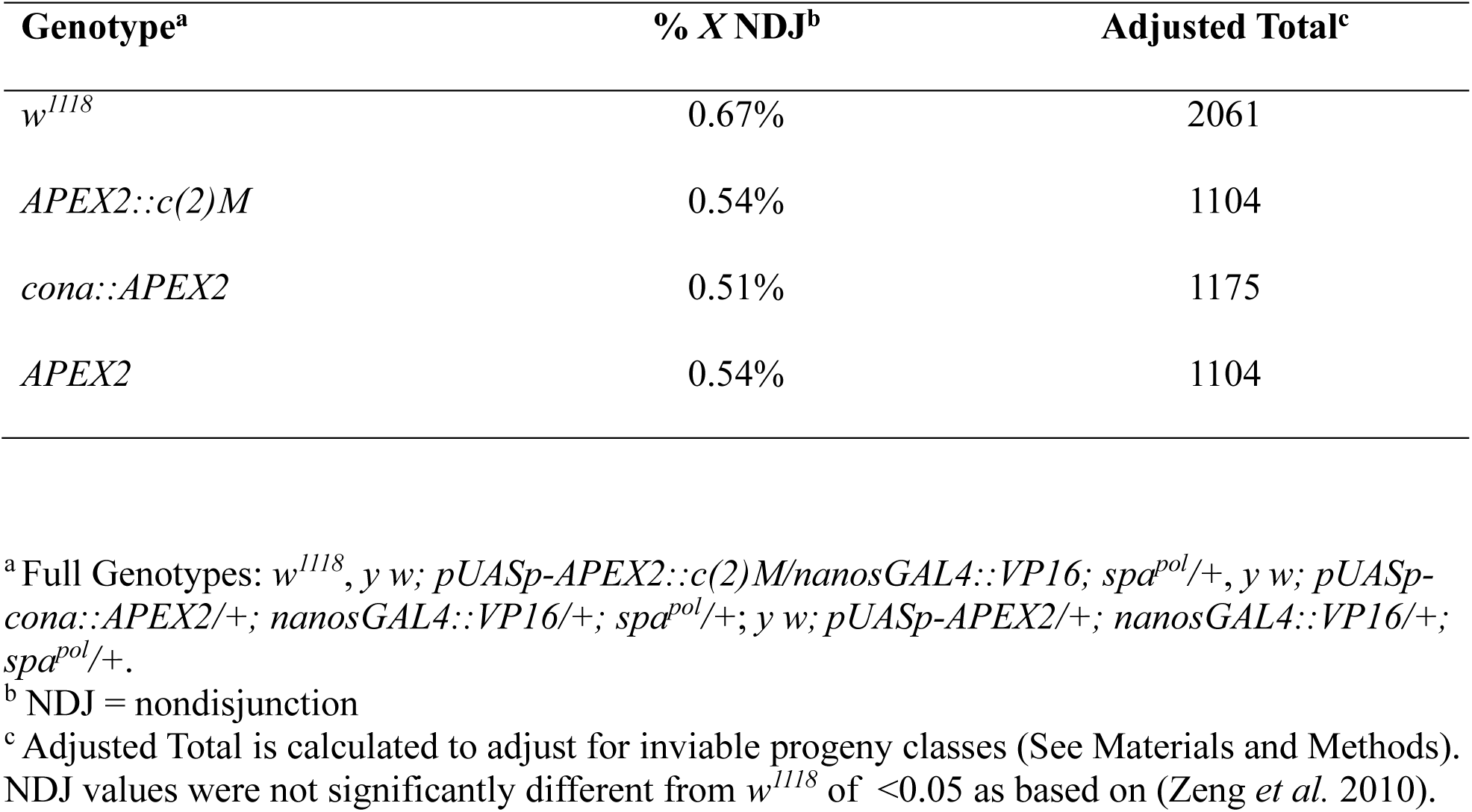
Expression of *APEX2* constructs do not affect chromosome segregation.

**Table 2.** Percent of *X* chromosome missegregation.

| Genotype<sup>a</sup> | %X NDJ<sup>b</sup> | Adj Total<sup>c</sup> | # Starting females<sup>d</sup> | Progeny per female |
| --- | --- | --- | --- | --- |
| Control | 0.2 | 1888 | 13 | 145.2 |
| *Cpsf5 RNAi* | 4.9** | 1154 | 15 | 76.9 |

<sup>a</sup> Control = *y w/y v; attP40/+; nanosGAL4::VP16/+; spa<sup>pol</sup>/+* and *Cpsf5 RNAi* = *y w/y sc v; TRiP.HMS00671)attP2 Cpsf5 RNAi/nanosGAL4; spa<sup>pol</sup>/+*.
<sup>b</sup> NDJ, nondisjunction
<sup>c</sup> Adjusted Total is calculated to account for inviable progeny classes (see Materials and methods).
<sup>d</sup> The number of female parents scored in the assays.
NDJ values between genotypes was statistically significant \*\* $p < 0.01$ as based on (Zeng *et al.* 2010).

APEX enzymes use biotin-phenol as a substrate with H2O2, to transfer biotin to suitable nearby electron-rich amino acids, such as Tyr, Trp, Cys, and His (Lam *et al*. 2015; Rhee *et al*. 2013). Biotin-phenol poses membrane-permeability challenges in living tissues which required the use of the detergent digitonin to successfully use APEX constructs to examine the ring canals of the *Drosophila* ovary (Mannix *et al*. 2019). Labeling conditions were modified from Mannix et al. (Mannix *et al*. 2019) (see Materials and Methods) to allow for maintenance of SC structures. The SC has been proposed to function as a biomolecular condensate (Von Diezmann *et al*. 2024) and the SC displayed sensitivity to the original detergent level. After biotin-labeling, normal SC was observed in the germaria by immunofluorescence (Fig. 2). By using streptavidin conjugated to Alexa-fluor 488 (Streptavadin-488), which binds to biotin, the extent of biotin-labeling could be evaluated by immunofluorescence to establish conditions with sufficient labeling (Fig. 2). Prior studies with tagged Cona constructs indicated that untagged Cona protein is preferentially built in full-length SC when both tagged and untagged versions are present (Cahoon *et al*. 2017). To improve incorporation of the Cona::APEX2 protein, the construct was used in a background where one endogenous copy of *cona* was mutated (*cona^f04903^/+)* (Page *et al*. 2008). Ovaries expressing either *APEX2::c(2)M*, *cona::APEX2*, or *APEX2* with the *nanosGAL4::VP16* driver were treated with biotin-phenol, along with *w^1118^* ovaries as a control for labeling by endogenous proteins (Fig. 2A-D). Comparing the localization of an antibody recognizing the transverse filament protein C(3)G with the streptavidin-488 showed that the streptavidin-488 was not observed to localize to any consistent subcellular compartment in the controls (Fig. 2A-B). In contrast, for both *cona::APEX2* and *APEX2::c(2)M* biotin-labeled ovaries, the streptavidin-488 could be observed to localize in a pattern similar to the SC in some nuclei (Fig. 2C-D). The apparent co-localization of streptavidin-488 to the SC suggests one or more proteins in or near the SC were labeled with biotin and that the Cona::APEX2 and APEX2::C(2)M proteins were incorporating into the SC. Not every SC positive nuclei showed the same level of biotin labeling, most likely due to variability in permeability of the labeling reagents and in the incorporation of the labeled APEX2 tagged proteins versions of the Cona and C(2)M proteins into the SC. The differences in localization of streptavidin-488 in the *cona::APEX2* and *APEX2::c(2)M* germaria compared to *APEX2* and *w^1118^* indicated that further analyses of the biotinylated proteins may yield proteins located in/near the SC.

The *nanosGAL4::VP16* driver promotes high expression of *pUASp* constructs in the female germline (Rorth 1998). Excess APEX tagged protein could be shuttled to locations other than the meiotic nuclei which would increase the number of biotin-labeled proteins without a role in meiosis. To enrich for meiotic targets, a Singulator 100 was used to isolate nuclei from biotin-labeled ovaries since the SC is associated with chromosomes. To isolate biotin-labeled proteins from isolated nuclei, nuclei were disrupted and applied to streptavidin-coated magnetic beads (See Material and methods). Biotinylated proteins isolated by streptavidin beads from nuclei of *APEX2*, *cona::APEX2*, *APEX2::c(2)M*, and *w^1118^* ovaries were sent for mass-spectrometry analysis in duplicate. The proteins from *APEX2::c(2)M*, *cona::APEX2*, and *w^1118^* samples were compared to the *APEX2* samples to provide a list of proteins enriched in those samples compared to the *APEX2* control sample. The *APEX2::c(2)M* samples had 173 proteins enriched compared to *APEX2* and the *cona-APEX2* showed enrichment for 180 proteins (Tables S1 and 2). Interestingly 65 proteins were in common between *APEX2::c(2)M* and *cona::APEX2* samples indicating these SC proteins were interacting with similar proteins or localized to similar locations. The *w^1118^* samples that underwent the same biotin-phenol labeling conditions showed enrichment for 22 proteins compared to the *APEX2* sample but none were in common with the *APEX2::c(2)M* and *cona::APEX2* enriched proteins (Table S3).

The proteins enriched in *APEX2::c(2)M* and *cona::APEX2* would be expected to be in close physical proximity to these proteins at some stage in the ovary since the distance APEX enzymes act is relatively small (Hung *et al*. 2016). The protein Nipped-B, which has been proposed to work in complex with C(2)M, was enriched in the *APEX2-c(2)M* and *cona::APEX2* samples (Gyuricza *et al*. 2016). The *cona::APEX2* samples showed an enrichment for SkpA, a member of the ubiquitin ligase complex SCF, which has been shown to be involved in regulating SC dynamics and karyosome morphology (Barbosa *et al*. 2021). Proteins with roles regulating transcription, for example Mediator complex components and su(var)3-3, the condensin subunit Cap-D2, and the centromere protein Cenp-C, were enriched in the *APEX2::c(2)M* and/or *cona::APEX2* samples. These proteins would be associated with the chromosomes, and therefore, could be in close proximity to SC components. Despite comparing the *APEX2::c(2)M* and *cona::APEX2* sample*s* to the *APEX2* control samples, there was enrichment for proteins associated with structures such as ribosomes and Golgi apparatus whose associations were likely found in the production and transport of the proteins. There were also proteins with only predicted functions or functions only described in somatic tissues. How these lesser-studied ovarian proteins potentially interact with the SC components in the female germline and whether they play biological roles in the germarium was of interest and the focus of this rest of the study. Proteins with published roles in female meiosis, the ovary, or basic cellular functions at the time of analysis were excluded from further study in favor of proteins with only predicted functions or with only limited information in non-ovarian tissues at the time of the analysis.

### RNAi knockdown of genes poorly characterized in the female germline revealed new information on protein functions

While some enriched proteins could associate with SC components before or after the SC is fully assembled, the immunofluorescence studies suggest that at least one or more proteins identified in the proteomics analysis are near or associated with fully assembled SC components. One of the lesser characterized enriched proteins could potentially play a role in meiotic processes such as SC assembly or DSB regulation in the female germline. We acquired available germline-expressed shRNAi hairpins targeting a subset of the *APEX2::c(2)M* and *cona::APEX2* enriched proteins from the Bloomington Stock Center that were created by the Harvard Transgenic RNAi Project (TRiP) (Zirin *et al*. 2020). A total of 80 lines targeting 63 genes were crossed to the *nanosGAL4::VP16* driver and examined for meiotic defects in the germarium (early to mid pachytene) using antibodies against the SC protein C(3)G and recognizing a phosphorylated form of histone 2A (γH2Av), a marker of DSBs (Table S4). As RNAi lines vary in the level of transcript knock-down, if multiple lines were available, all were examined.

In wild-type females in region 2A (early pachytene) of the germaria, up to four nuclei of each 16-cell cyst build full-length track-like SC, and DSBs are induced, which can be visualized using an antibody recognizing γH2Av. In region 2B (early-mid pachytene) SC disassembly begins in the nuclei not selected to be the oocyte nucleus and γH2Av foci decrease in number as DSB repair is underway. By region 3 (mid-pachytene) only the selected oocyte nucleus has full-length SC and γH2Av foci are absent. For scoring the shRNAi lines, a defect in SC assembly was assigned for lines failing to form full-length SC in multiple nuclei within region 2A (early-pachytene). A defect in SC maintenance was assigned for lines failing to retain a nucleus with full-length SC at region 3 (mid-pachytene). For DSB phenotypes, germaria were scored for the absence of γH2Av foci in region 2A, which would indicate a failure in DSB induction, or retention of γH2Av foci in region 3, which indicates a DSB repair defect.

For five of the shRNAi lines tested, expression with *nanosGAL4::VP16* led to mostly underdeveloped ovaries, with few, if any later stage oocytes, indicating strong defects in the mitotic divisions of germline stem cells, meiotic entry, or ovarian development (Table S4). Three lines displayed defects in SC dynamics. Compared to control germaria (Fig. 3A), germaria expressing an RNAi construct targeting *NOP2-Sun domain family member 2* (*Nsun2*, *CG6133*, BL62495) displayed a reduction in the number of nuclei with SC assembled in region 2A, but nuclei with full-length SC could be observed in regions of the germaria typically corresponding to the late 2B and 3 regions (Fig. 3B, Table S). The delay in SC formation was accompanied by delays in or absence of γH2Av foci. Since full-length SC was eventually formed it suggests there are delays either in oocyte development and/or meiotic entry upon *Nsun2* RNAi knock-down.

**Figure 3.**
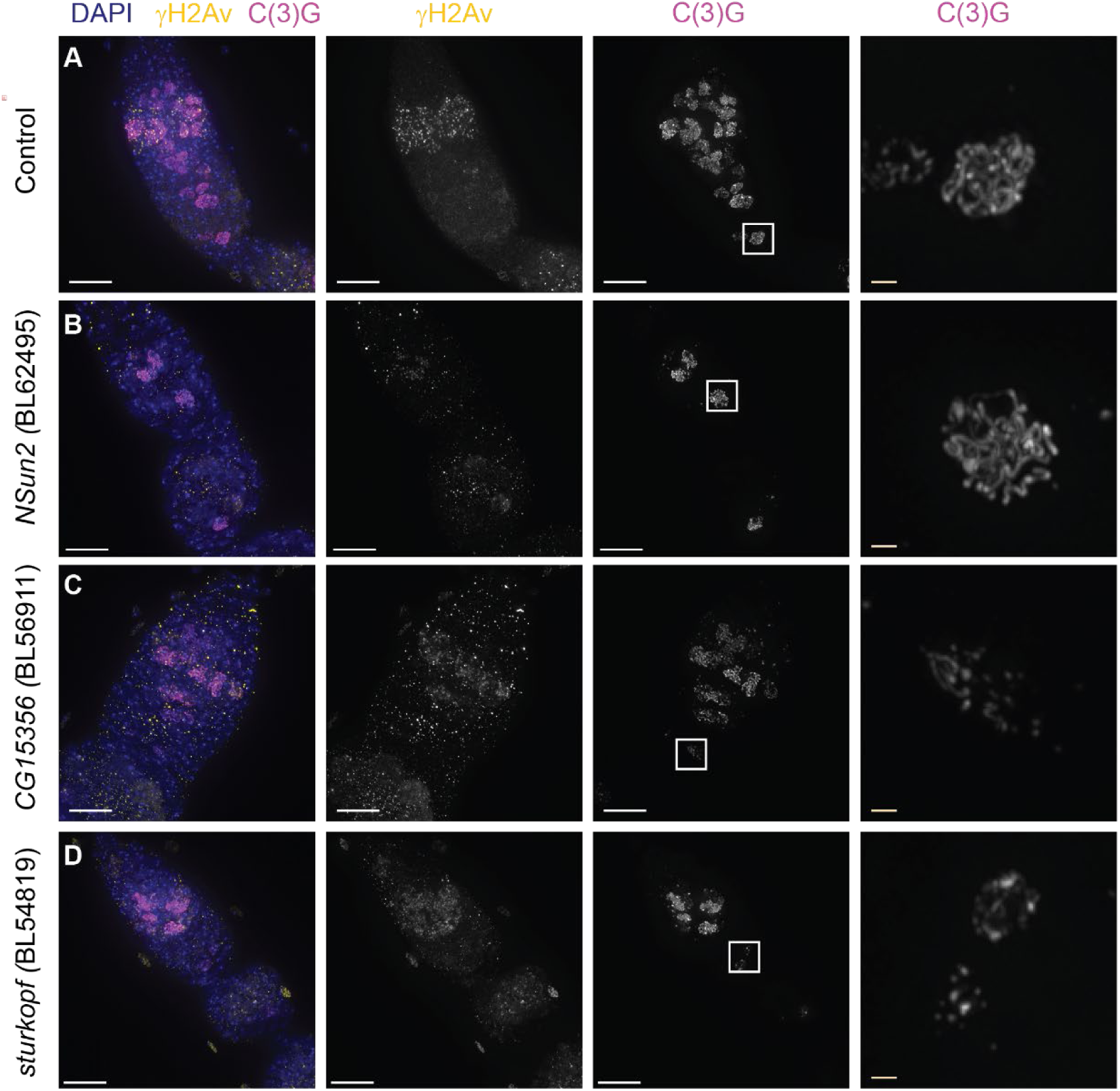
Germline knock-down of genes enriched in proximity labeling cause defects in SC dynamics. (A) Control (*w^1118^*) germarium and region 3 nucleus displaying normal γH2Av (DBS – yellow) and SC (C(3)G antibody -magenta) dynamics. (B) Germline knock-down of *Nsun2* (BL62495) causes a delay/reduction in nuclei with full-length SC. As γH2Av dynamics were also altered it suggests developmental delays. Approximate region 2B nucleus is shown magnified. (C-D) Knock-down of *CG15356* (BL56911) (C) and *sturkopf* (BL54819) (D) causes early disassembly of the SC. Magnified nuclei are from approximately region 3 (C) and late region 2B (D). γH2Av foci appear and disappear with normal dynamics indicating the defect is SC specific. RNAi lines were driven with *nanosGAL4::VP16*. Boxed regions are shown enlarged in the last column. Scale bar = 10 µm and inset = 1µm. Images are projections from z-stacks.

The shRNAi lines targeting *sturkopf* (*CG9186*, BL54819) (Fig. 3C) and *CG15356* (BL56911) (Fig. 3D) caused a defect in SC maintenance, with the region 3 nucleus displaying some evidence of partial SC fragmentation (Table S4). Sturkopf plays a role in lipid droplet formation (Thiel *et al*. 2013) and *CG15356* was reported to associate with the SIN3 histone deacetylase protein in S2 cells (Spain *et al*. 2010). In both lines, γH2Av foci appeared in region 2A and were absent from region 3 (Fig. 3C-D) suggesting the knock-down did not affect DSB induction or repair. As the knowledge of the function of these genes in other tissues/cell types fail to suggest the mechanism of the observed defects further work is needed to determine the specific function in the germarium of these proteins and how they may interact with SC proteins.

### A shRNA line targeting a predicted member of a mRNA cleavage factor complex leads to multiple meiotic defects

RNAi knock-down targeting the gene *Cleavage and polyadenylation specific factor 5* (*Cpsf5*) (BL32883) caused defects in SC assembly. The Cpsf5 protein is predicted to be involved in mRNA processing as part of an mRNA cleavage factor complex. *Cpsf5* RNAi expressing germaria displayed a range in the severity of the SC phenotypes compared to control germaria (Fig 4.A-D). In the most severely affected *Cpsf5 RNAi* germaria, nuclei with only puncta of SC (zygotene-like) or partial SC tracks of SC were observed (Fig. 4B), while control germaria displayed many nuclei with full-length SC throughout all three meiotic stages of the germaria (Fig. 4A). In other *Cpsf5 RNAi* germaria, nuclei with mostly full-length or full-length SC were observed but in only some regions of the germaria and the total number of nuclei with full-length SC was reduced (Fig. 4C-D). The variability in the severity of the SC phenotype between *Cpsf5 RNAi* germaria may be due to variation in the level of RNAi knockdown between germaria.

**Figure 4.**
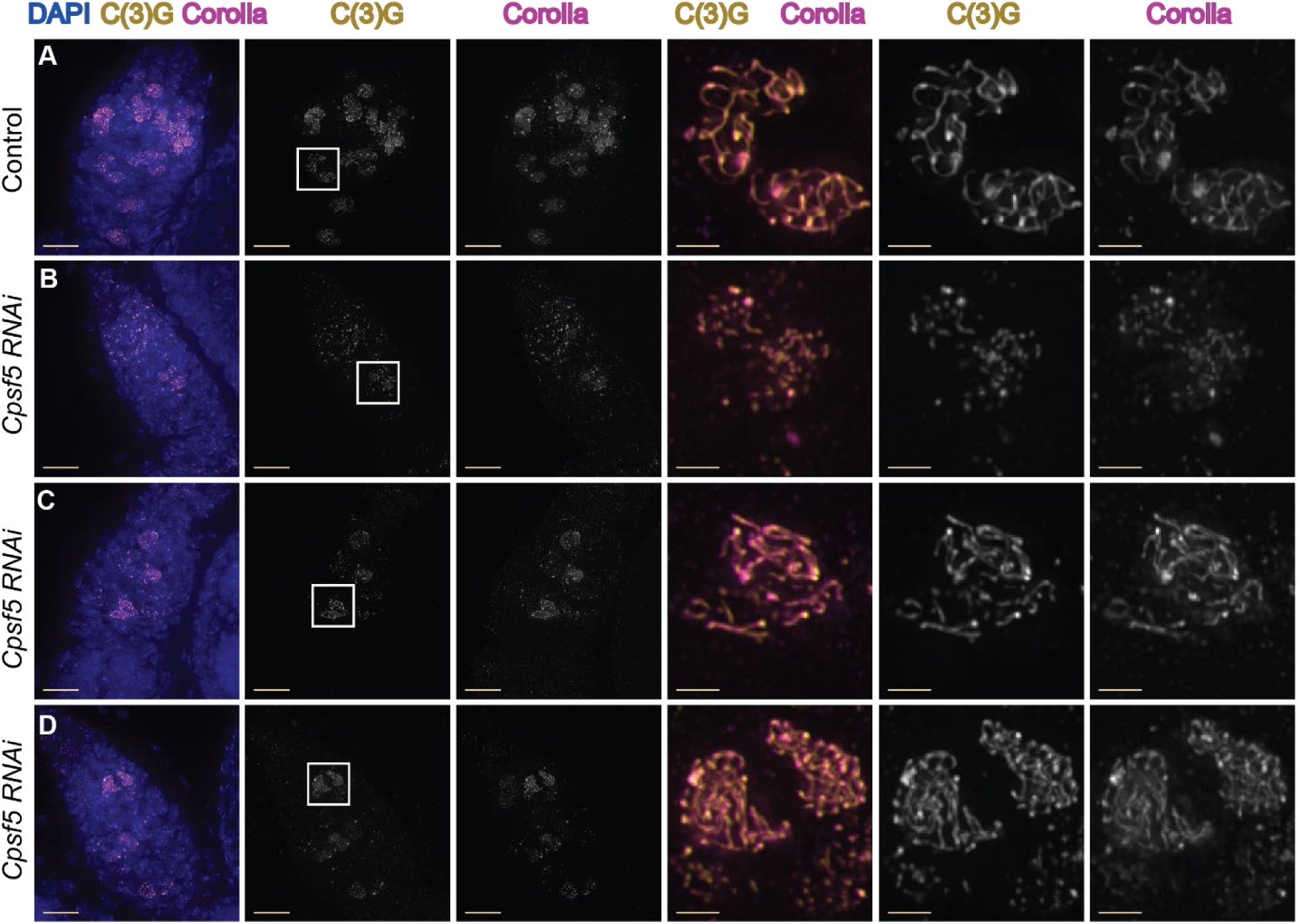
Germline knock-down of *Cpsf5* driven by *nanosGAL4::VP16* causes defects in SC assembly. (A-D) Shown are antibodies recognizing the central region SC components C(3)G (yellow) and Corolla (magenta) with DAPI to visualize DNA (blue). (A) Control germarium (*attp40/+; nanosGAL::VP16/+*) shows long tracks of both SC components in multiple regions of the germarium. (B-D) Germaria expressing *Cpsf5 RNAi* (BL32883) by *nanosGAL4::VP16* form only puncta/short tracks of SC components (B) or only partial/full length SC in a few nuclei (C-D). Full genotype is *y^1^ sc* v^1^ sev^2^/ y w ^1^; P(y^+t7.7^ v^+t1.8]=TRiP.HMS00671^ attP2 /nanosGAL4::VP16; spa^pol^/+*. Scale bars = 10 µm, inset 2=µm. Images are projections from z-stacks.

**Fig. 5.**
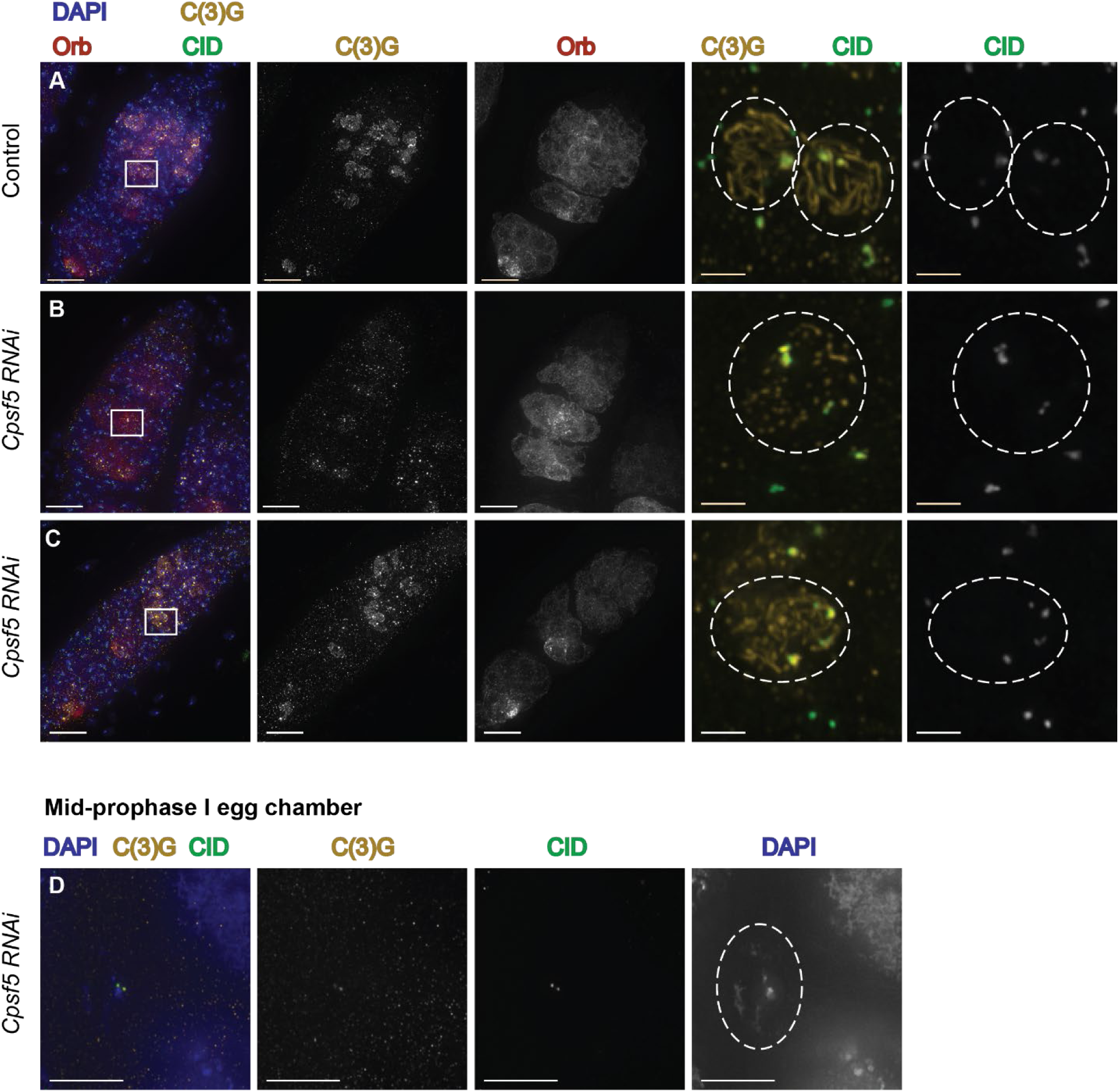
*Cpsf5 RNAi* ovaries display modest defects in centromere clustering. (A-D) Antibodies recognizing CID (centromeres, green), C(3)G (SC, gold), and Orb (oocyte development, dark red) are shown with DAPI (DNA, blue). Boxes indicate the areas shown magnified to the right. (A) Control (*attp40/+; nanosGAL4::VP16/+*) germarium shows normal full-length SC formation and magnified box shows nuclei with one and two centromere clusters. (B-C) Show examples from *Cpsf5 RNAi* germaria (*y^1^ sc* v^1^ sev^2^/ y w ^1^; P(y^+t7.7^ v^+t1.8]=TRiP.HMS00671^ attP2 /nanosGAL4::VP16; spa^pol^/+*). (B) Germarium displays mostly punctate SC while centromeres are clustered in two regions. Orb is expressed in the cysts of region 2A and begins to accumulate around a single nucleus as the cysts progress. (C) Germarium with several nuclei with mostly full-length SC. Orb expression is initially weak but does show some accumulations around the pro-oocytes in later regions. Magnified region shows a centromere clustering defect despite the presence of SC tracks. (D) Example of a *Cpsf5 RNAi* mid-prophase oocyte with a karysome defect. Despite the defect, C(3)G is still associated with the centromeres. Scale bar = 10 µm, magnified region = 2 µm. Images are projections from z-stacks.

With strong defects/delays in full-length SC assembly, we examined the *Cpsf5 RNAi* females for additional meiotic defects. In *D. melanogaster* females, loss of SC components leads to chromosome missegregation due to a failure in crossover formation (Collins *et al*. 2014; Page and Hawley 2001; Page *et al*. 2008). Segregation of the *X* chromosome was examined in *Cpsf5 RNAi* females compared to *nanosGAL4::VP16* control flies. A modest, but statistically significant, increase in *X* chromosome nondisjunction of 4.9% was observed. Additionally, the genetic assay revealed a decrease in fertility of the *Cpsf5 RNAi* females with approximately half the progeny produced per female compared to control females.

We next examined centromere clustering, as SC components are required to cluster centromeres into an average of two centromere clusters during pachytene (Collins *et al*. 2014; Takeo *et al*. 2011). Centromere clustering was scored using an antibody to Centromere Identifier (CID), the Cenp-A homolog in *Drosophila,* a C(3)G antibody to mark SC, and an antibody to Orb to help identify the pro-oocyte nucleus in regions late 2B and 3, even if SC was fragmented. A statistically significant increase in the average number of CID clusters was found for *Cpsf5 RNAi* ovaries compared to control ovaries for region 2A, 2B and mid-prophase stages (Table 3). Nuclei in region 3 of *Cpsf5 RNAi* germaria also displayed an increase in the average number of CID foci but due to the smaller number of nuclei that could be scored the result did not reach statistical significance (p=0.0756) (Table 3). Figure 5A shows a control germarium with a nucleus with the expected one or two CID clusters by immunofluorescence. Figure 5B-C shows examples from *Cpsf5 RNAi* ovaries where centromeres have clustered and where centromere clustering is defective, respectively. The centromere clustering defect in *Cpsf5 RNAi* females is less severe than that observed in null mutants of SC components (Collins *et al*. 2014; Takeo *et al*. 2011). Additionally, the strength of SC defect based on immunofluorescence was not always correlated with the strength of centromere defect in individual nuclei (Fig. 5B-C). For the centromere clustering assay mid-prophase egg chambers were scored. This *Cpsf5 RNAi* line (BL32883) was identified previously in a screen for karyosome defects in mid-oogenesis, but the line was not examined for further defects (Nieken *et al*. 2023). Karyosome defects were observed in some of the mid-prophase egg chambers scored for centromere clustering (discussed below) (Fig. 5D). The defects in centromere clustering supports that *Cpsf5 RNAi* knock-down is affecting a function of the SC.

**Table 3.** Centromere clustering (CID foci) by meiotic stage

| Genotype<sup>a</sup> | Region 2A<br>(Early-Pachytene) | Region 2B<br>(Early/mid-pachytene) | Region 3<br>(Mid-pachytene) | Stages 2-8<br>(Mid-prophase) |
| --- | --- | --- | --- | --- |
| Control | 1.5 +/-0.6<br>N=48 | 1.7 +/-0.5<br>N=26 | 1.8 +/-0.7<br>N=15 | 1.9 +/-0.7<br>N=33 |
| *Cpsf5 RNAi* | 2.3 +/-0.9**<br>N=16 | 2.6 +/-0.9**<br>N=17 | 2.4 +/-0.9<br>N=9 | 2.5 +/-1.2*<br>N=31 |

<sup>a</sup> Control = *y w/y v; attP40/+; nanosGAL::VP16/+; spa<sup>pol</sup>/+* and *Cpsf5 RNAi* = *y w/y sc v; TRiP.HMS00671)attP2 Cpsf5 RNAi/nanosGAL4::VP16; spa<sup>pol</sup>/+*. P values by Mann-Whitney U test \* $<0.5$ and \*\* $<0.01$ .

To further investigate the effect of *Cpsf5 RNAi* knock-down on SC function, we examined crossing over on the left arm of the *2^nd^* chromosome. In *D. melanogaster* females, null mutations of SC components have severe defects in crossover formation and mutations that partially perturb the SC display changes in crossover placement and reductions in crossing over in some genetic intervals (Billmyre *et al*. 2019; Williams *et al*. 2025). The effect of *Csf5 RNAi* on crossing over on the *2^nd^* chromosome was examined from the tip of *2L* to a marker across the centromere on *2R* (Table 4). *Cpsf5 RNAi* females displayed a reduction in total crossing over and an increase of non-crossover chromatids (65.3% compared to 51.1% for the control). This increase of non-crossover chromatids was offset by decreases in the recovery of single crossover chromatids (31.9% compared to 45.3% for the control). As only one of the four chromatids can be recovered during female meiosis, the rate of crossovers must be mathematically derived using Weinstein tetrad analysis with the expected rate of non-crossover, single crossover, and double crossovers being designated E0, E1, E2 respectively (Weinstein 1936; Zwick *et al*. 1999). The E0 Weinstein tetrad value for *Cpsf5 RNAi* females was increased (0.3591) compared to control females (0.0935) while the E1 value was decreased (0.4659 for *Cpsf5 RNAi* females compared to 0.7647 for controls). RNAi knockdown of *Cpsf5* leads to a decrease in total recombination for the *2^nd^* chromosome.

**Table 4.**
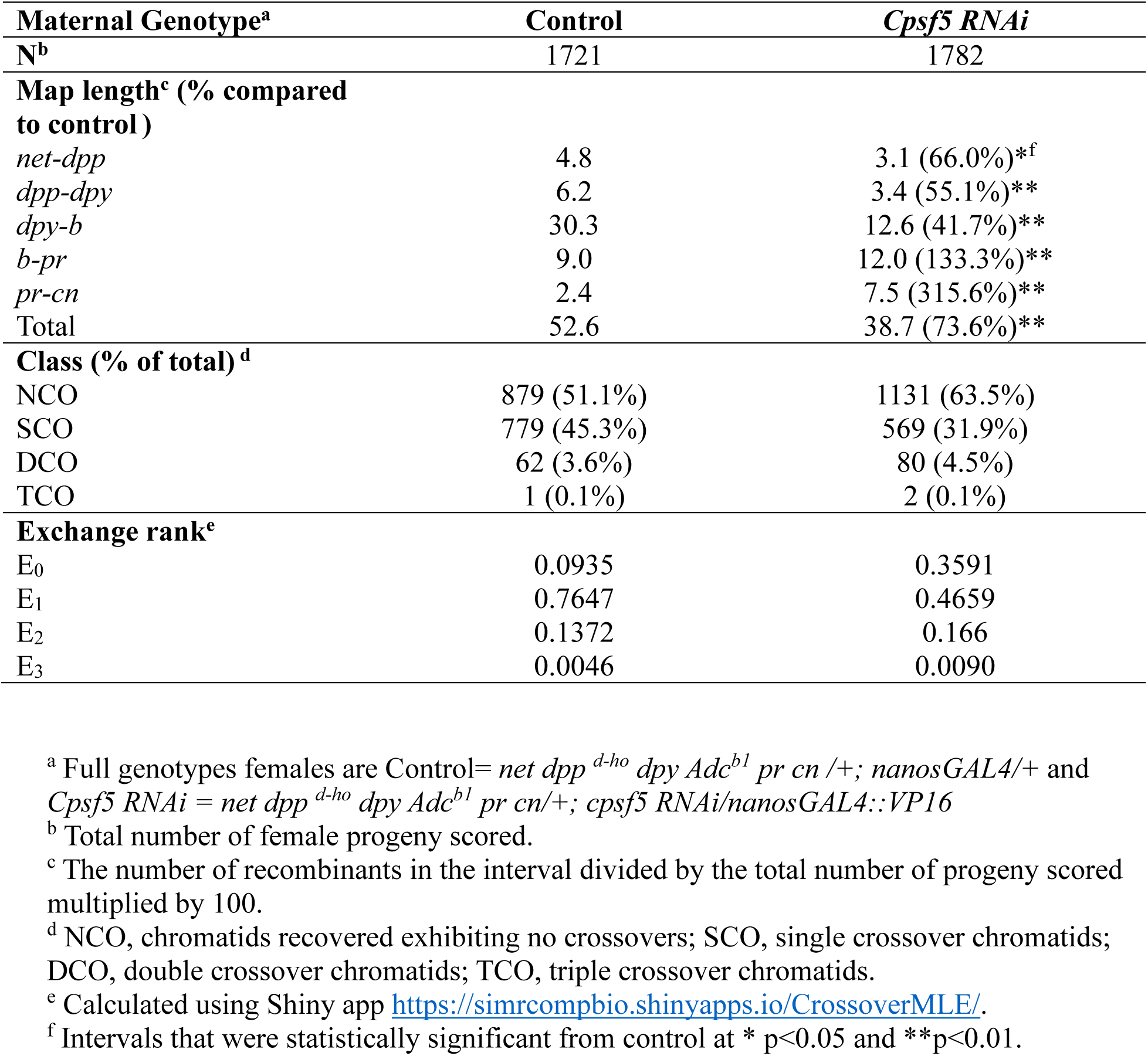
Placement of crossovers are affected on the *2^nd^* chromosome in *Cpsf5 RNAi* females.

### RNAi knock-down of *cpsf5* caused a retention of DSBs and a change in crossover placement on an autosome

As mutations in SC components in *D. melanogaster* females cause changes in crossover placement (Billmyre *et al*. 2019; Hawley *et al*. 2025; Williams *et al*. 2025), we examined the level of crossing over within each genetic interval. The *cpsf5 RNAi* females showed statistically significant decreases in crossing over in the genetic intervals distal from the centromere (*net-b*) but a strong increase in crossing over compared to the control (315.6% increase) in the interval spanning the centromere of chromosome *2* (*pr*-*cn*). A modest increase was also observed in the *b*-*pr* interval. *Cpsf5* knock-down females displayed both a decrease in total recombination on the *2^nd^* chromosome and a shift in crossover placement towards the centromere.

In wild-type flies, γH2Av foci, which marks DSBs, are rarely observed at region 3, due to the repair of DSBs by the meiotic repair machinery. During the initial screening of the *Cpsf5 RNAi* line, the presence of γH2Av foci was observed at region 3 (mid-pachytene), indicating defects in DSB repair in *Cpsf5* knock-down germaria. To better characterize the extent of the repair defects, germaria from *Cpsf5 RNAi* and control females were analyzed with antibodies to C(3)G to mark SC, γH2Av to mark DSBs, and Orb to help identify the oocyte at regions 2A, 2B, and 3. In control females, numerous γH2Av foci could be observed in region 2A with the number of foci decreasing until 0 foci were observed in the Orb selected region 3 nuclei (Table 5, Figure 6A). The *Cpsf5* knock-down germaria displayed a similar average number of γH2Av foci compared to the control in region 2A (early-pachytene) (Table 5, Fig. 6B-E) suggesting DSBs are being initiated at wild-type levels in most *Cpsf5 RNAi* germaria. In region 2B (early-mid pachytene), while the control displayed a strong decrease in the average number of γH2Av foci (0.6 average), only a small average decrease was observed in the *Cpsf5 RNAi* germaria (9.2 average γH2Av foci). By region 3 (mid-pachytene), γH2Av foci were still prevalent (average 7.1 foci γH2Av foci) in the *Cpsf5 RNAi* germaria. Shown are examples at the perdurance of γH2Av foci in region 3 nucleus with partial SC (Fig 6B), punctate SC (Fig 6C) and mostly full-length SC (6D). Examples of germaria with more wild-type γH2Av dynamics were observed in a few *Cpsf5* knock-down ovaries (Fig 6E) suggesting DSB repair may be occurring in some germaria. The observation of γH2Av foci in region 3 indicates a defect in DSB repair in *Cpsf5 RNAi* germaria.

**Fig. 6.**
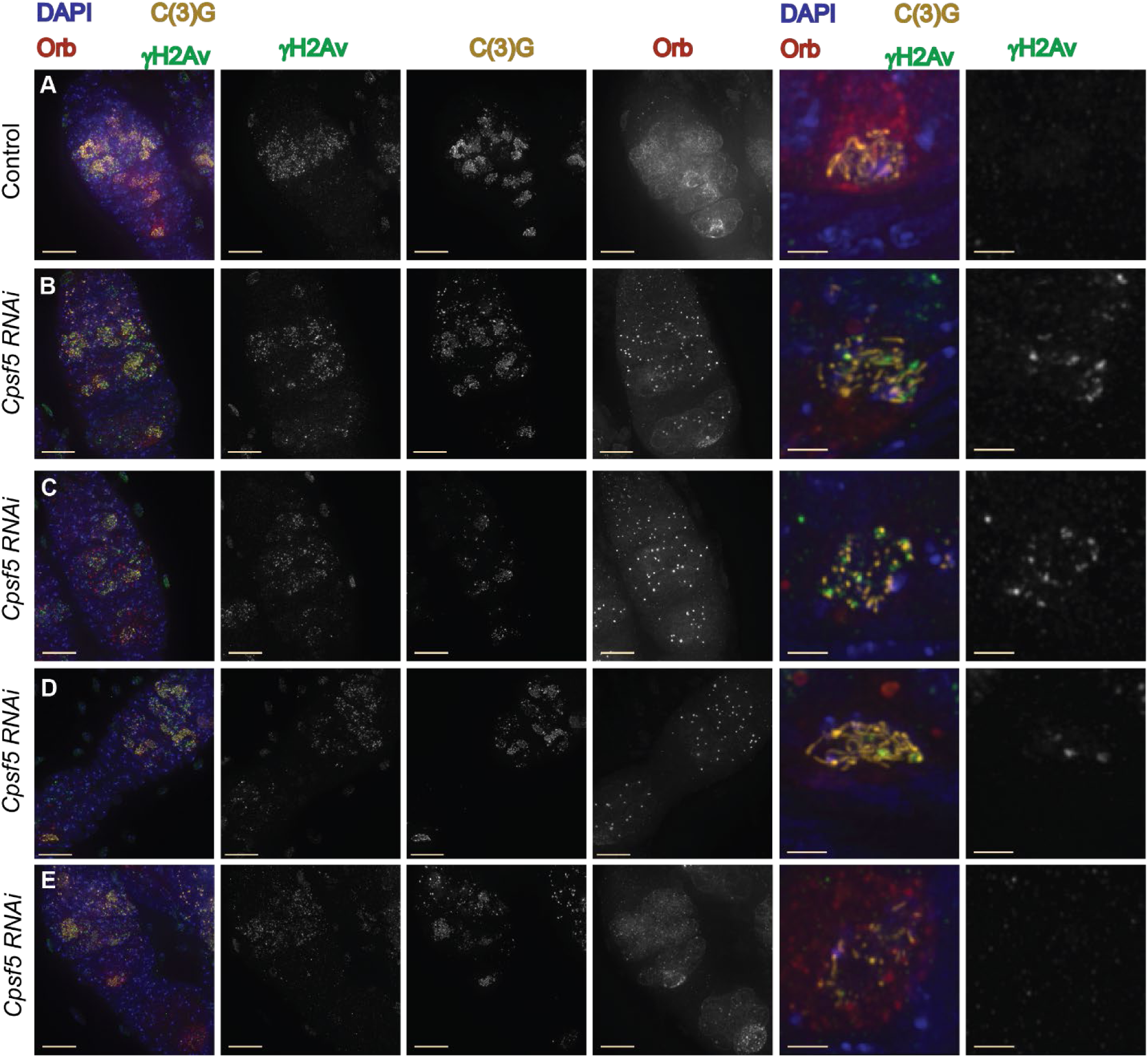
γH2Av foci persist in *cpsf5 RNAi* germaria. (A-E) Shown are germaria labeled with antibodies to γH2Av (DSBs, green), C(3)G (SC, gold), Orb (oocyte development, dark red), and DAPI (DNA, blue). Boxed nuclei show enlarged. (A) In the control (*y w/ y v; attP40/+; nanosGAL::VP16/+; spa^pol^/+)* germarium many γH2Av foci can be observed in region 2A where numerous nuclei with full-length SC are present. γH2Av foci decrease in number through the later stages of the germarium with no foci present in the region 3 oocyte nuclei (boxed region shown magnified). (B-E) Defects in γH2Av dynamics are observed in *Cpsf5 RNAi (y^1^ sc* v^1^ sev^2^/ y w ^1^; P(y^+t7.7^ v^+t1.8]=TRiP.HMS00671^ attP2 /nanosGAL4::VP16; spa^pol^/+)* germaria. (B) γH2Av foci are present in region 2A but persist in the Orb selected oocyte nucleus with partial tracks of SC in region 3. (C) γH2Av foci are present in region 2A but Orb expression is weak and γH2Av foci persist in the nucleus with punctate SC in region 3. (D) While more nuclei formed full-length SC, γH2Av foci still persist in the region 3 nucleus and Orb localization is aberrant. (E) Germarium where γH2Av foci are absent from the Orb selected pro-oocyte nucleus in region 3, but the SC is not full length. Scale bar = 10 µm, magnified region = 2 µm. Images are projections from z-stacks.

**Table 5.** γH2Av foci persist into Region 3 (Mid-pachytene) in *Cpsf5 RNAi* females

| Genotype<sup>a</sup> | Region 2A (Early-Pachytene) | Region 2B (Early/mid-pachytene) | Region 3 (Mid-pachytene) |
| --- | --- | --- | --- |
| Control | 12.3 (+/- 4.7)<sup>b</sup> | 0.6 (+/-1.2) | 0 (+/-0) |
|  | N<sup>c</sup>=32 | N=29 | N=17 |
| *Cpsf5 RNAi* | 11.6 (+/-3.5) | 9.2 (+/-4.8)** | 7.1 (+/-5.3)** |
|  | N=26 | N=13 | N=14 |

<sup>a</sup> Control = *y w/ y v; attP40/+; nanosGAL::VP16/+; spa<sup>pol</sup>/+* and *Cpsf5 RNAi* = *y w/ y sc v; TRiP.HMS00671)attP2 Cpsf5 RNAi/nanosGAL4; spa<sup>pol</sup>/+*.
<sup>b</sup> Average number of γH2Av foci with standard deviation.
<sup>c</sup> N=total number of nuclei scored.
\*\* Statistically significant from the control at the same stage at <0.01.

The persistence of DBSs in region 3 can induce later oocyte phenotypes, including karyosome defects and sterility (Ghabrial *et al*. 1998). The γH2Av foci observed in region 3 provides an explanation for the previously observed karyosome defect in mid-prophase *Cpsf5 RNAi* egg chambers (Nieken *et al*. 2023). During the analysis of centromere clustering in mid-prophase some normal spherical karyosomes were observed. These may be derived from the subset of germaria with few or no γH2Av foci in region 3 nuclei. The nondisjunction assays revealed an approximately 50% reduction in fertility for *Cpsf5 RNAi* females. It seems likely that decreased female fertility is caused by those egg chambers with the most severe defects in DSB repair and karyosome formation undergoing arrest/ death. The germaria and egg chambers with more wild-type DSB dynamics and karyosome formation in *Cpsf5 RNAi* ovaries likely give rise to progeny recovered in genetic assays.

### In *Cpsf5* knock-down germaria, the Orb protein showed variability in accumulation suggesting defects in the coordination of oocyte development and meiotic events

Orb is a marker of cyst development and oocyte specification in the *Drosophila* female germline, with Orb expression in the cytoplasm of the entire cyst after meiotic entry and Orb progressively accumulating around the nucleus selected to be the oocyte nucleus (Fig 5A and 6A) (Lantz *et al*. 1992; Lantz *et al*. 1994; Lantz and Schedl 1994). Orb expression was variable both between cytological preparations and within a cytological preparation of *Cpsf5 RNAi* ovaries (Fig 5B-C, 6B-E). Some germaria displayed a more wild-type Orb protein localization pattern (Figure 5B). In other germaria, the diffuse Orb protein localization appeared weaker in the cysts of region 2A and 2B (Fig 5C, 6C-D) and/ or Orb failed to strongly accumulate around the pro-oocyte in late region 2B and region 3 (Fig 6C-D). Some of the variability may be due to variation in the level of RNAi knock-down of *Cpsf5* expression. The degree of severity of Orb defects did not always correlate with the level of SC, DSB, or centromere clustering defect. For example, Fig. 5B shows a stronger SC defect but more robust Orb expression than Fig. 5C or 6D. Retention of γH2Av foci in region 3 could be observed in germaria both with accumulation of Orb antibody around the pro-oocyte (Fig. 6B) and in examples with weak or absent Orb accumulation (Fig. 6C-D).

We found that *Cpsf5 RNAi* germaria display defects in SC formation, a persistence of γH2Av foci, chromosome missegregation, and decreases in fertility. The myriad of meiotic defects suggests Cpsf5, and potentially RNA processing in general, plays a role in coordinating and promoting proper meiotic progression in the female germline. Interestingly, overexpression of *Cpsf5*, but not RNAi knock-down, caused defects in the switch from mitosis to meiosis in the *Drosophila* male germline (Shan *et al*. 2017). Evidence suggests the *Cpsf5* transcript is repressed by the Tut-Bam-Bgcn proteins that promotes the mitosis-to-meiosis transition in males (Shan *et al*. 2017). The results of the RNAi knock-down of *Cpsf5* in the female germline and the repression of *Cpsf5* in the male germline suggests Cpsf5 and 3’RNA processing may be involved in promoting female meiotic progression. It would be interesting in future studies to investigate if *Cpsf5 RNAi* alters the 3’UTR patterns of RNA transcripts and to compare any changes to those observed in the male germline after manipulation of the RNA 3’ -processing machinery (Shan *et al*. 2017). How the Cpsf5 protein is associated with the Cona protein during female meiosis will also require further investigation. Cpsf5 is predicted to part of the machinery regulating the alternative polyadenylation of gene transcripts (Shan *et al*. 2017; Tang *et al*. 2018). Changes in 3’UTR length are associated with changes in the localization, stability and translation efficiency of mRNAs (reviewed in (Tian and Manley 2017)). One possibility is that Cpsf5 is associated with Cona to help coordinate the timing of meiotic events with the appearance of SC proteins.

### Proximity ligation revealed a new regulator of meiotic events in the female germline

Despite decades of investigation, many aspects of meiosis are still poorly understood at the molecular level. While earlier methods relied on genetic screens to identify proteins involved in *Drosophila* female meiosis, this approach tends to yield only certain classes of proteins. To identify additional proteins with roles in the early *Drosophila* female germline, we adapted the proximity labeling approach which had been used successfully to investigate protein interactions in other *Drosophila* processes (Chen *et al*. 2015; Mannix *et al*. 2019). The goal was to identify proteins with close physical proximity to known SC proteins using APEX2 tagged proteins with the prospect that a subset of those proteins would also play roles in female meiosis. Biotinylated proteins found enriched in the *APEX2::c(2)M* and *cona::APEX2* samples compared to controls by mass spectrometry included proteins expected to localize with the homologous chromosomes, but also proteins less characterized in the early female ovary. Examining available RNAi lines targeting a subset of these genes revealed lines with ovarian and meiotic phenotypes, providing preliminary evidence for functions in the early female germline for these proteins (Table S4). A line targeting *Cpsf5,* implicated in 3’ RNA processing (Tang *et al*. 2018) caused defects in multiple meiotic processes, implicating 3’ RNA processing as a mechanism of meiotic regulation, including SC assembly, in the female germline. As *Cpsf5* mutants die as larvae, a role for Cpsf5 in early female meiosis would not have been revealed through traditional forward genetic screens (Tang *et al*. 2018). Our proximity labeling experiment revealed a potential new pathway of study for the regulation of *Drosophila* female meiosis and illustrates the value of using a proximity labeling approach to reveal new proteins with roles in meiosis. Future proximity ligation studies with additional meiotic proteins, and possibly with alternative enzymes (reviewed (Samavarchi-Tehrani *et al*. 2020)), have the potential to yield additional novel meiotic proteins.

## Materials and Methods

### Drosophila husbandry

Flies were maintained at 25°C on standard food. The germline-specific *nanosGAL4::VP16* driver (*y w; Pnanos-GAL4::VP16; spa^pol^*) was used to express both the APEX2 tagged constructs and the UAS-RNAi lines (Rorth 1998). For proximity ligation experiments, the genotypes *y w; pUASp-APEX2 (attP40)*, *y w; pUASp-APEX2::c(2)M (attP2)*, or *y w; pUASp-cona::APEX2 (attP40); cona^f04903^/TM3* were crossed to the *nanos-GAL4::VP16* driver. *w^1118^* was used as the no construct control. All *Drosophila* RNAi stocks tested were created by the Harvard Transgenic RNAi project (TRiP collection) (Zirin *et al*. 2020) and acquired from the Bloomington *Drosophila* Stock Center (BDSC) prior to September 2024 (Table S4). Only TRiP lines constructed in the germline expressed VALIUM20 or VALIUM22 vectors were chosen. While the VALIUM 21 drives germline expression, no RNAi lines targeting a gene of interest was available in this vector at the time of screening. See Table S4 for a list of stocks from the Bloomington stock center tested for the RNAi screen. Full information for these lines can be found on the TRiP website (https://fgr.hms.harvard.edu/trip-in-vivo-fly-rnai). Controls for the RNAi screen were *w^1118^*, *y w; spa^pol^*, or females generated from crossing Bloomington lines BL36304 *y^1^ v^1^; P(y+t7.7]=CaryP)Msp300[attP40]* to *y w; nanosGAL4; sv^spa^-^pol^*. Genotype of RNAi knock-down line targeting *Cpsf5* was BL32883 *y^1^ sc* v^1^ sev^21^; P(y^+t7.7^ v^+t1.8]=TRiP.HMS00671^ attP2*.

Gene names provided in the RNAi screen Table S4 were those listed in FlyBase (https://flybase.org) as of June 2025 (Gramates *et al*. 2022). Short descriptions of the genes provided in Table S4 were acquired from FlyBase and may not cover all functions predicted for the genes (Gramates *et al*. 2022).

### Construction of the APEX2 constructs

To construct the *UASp*-driven *APEX2::c(2)M* transgene, the *c(2)M* coding region with stop codon was PCR amplified from a cDNA containing plasmid with NdeI and XbaI restriction sites. *APEX2* was PCR amplified from pcDNA3 APEX2-NES (Addgene Plasmid #49386) without both the stop codon and nuclear export signal with NotI and NdeI restriction sites. The *pUASp-attB* vector (NotIHF and XbaI), the *APEX2* fragment (NotI and NdeI), and the *c(2)M* fragment (NdeI and XbaI) were digested and gel purified. Both fragments were subcloned together into the *pUASp-attB* vector.

For the *pUASp*-driven *cona::APEX2* transgene, the coding region of *cona* without stop codon was PCR amplified from a vectoring containing the *cona* cDNA with NotI and NdeI restriction sites. *APEX2* with stop codon but nuclear export signal removed was PCR amplified with NdeI with XbaI restriction sites. The *pUASp-attB* vector (NotIHF and XbaI), the *cona* (NotI and NdeI), and the *APEX2* (NdeI and XbaI) fragments were digested and gel purified. Both fragments were subcloned together into *pUASp-attB*.

For the *pUASp*-driven *APEX2* transgene, the coding region of *APEX2* with stop codon but without the nuclear export signal was PCR amplified with NotI and XbaI restriction sites. The *pUASp-attB* vector and the APEX2 fragment were digested with NotIHF and XbaI and gel purified. The APEX2 fragment was subcloned together into *pUASp-attB*.

All three constructs were sequence verified and sent to GenetiVision Corporation for injection using ϕC31 site-specific integration into the *attP40* (*APEX2* and *APEX2::cona*) or *attP2* (*APEX2::c(2)M*) lines.

### Biotin labeling

Females that were 1-3 days post-eclosion were provided wet yeast and males overnight. Up to 34 pairs of ovaries were dissected in PBS pH 8 and placed in a 1.5 mL tube. Tubes were spun down at low speed to allow removal of residual PBS. 500 ul of a 0.165% digitonin and 0.5 mM Biotin phenol in PBS pH.8 solution was added to the ovaries and the ovaries were rocked at room temperature for 10 minutes. H2O2 (hydrogen peroxide) was freshly diluted to 100 mM in PBS pH 8 and 2 µL was applied to the ovary solution for 1 minute while shaking. 5 µL of 1 M NaH3 (Sodium Azide) was added to quench the reaction. Ovaries were allowed to quickly settle, and the solution was removed. Ovaries were washed rapidly 3 times with mixing with 10 mM sodium Azide solution diluted in PBS pH 8. Labeled ovarian samples were either flash-frozen in liquid nitrogen and stored at -70°C or promptly fixed for cytology (see Immunofluorescence).

### Isolation of biotinylated proteins

For isolation of biotinylated proteins, biotin-phenol treated ovary samples were combined to a total of approximately 100 ovary pairs in 300 µl cold PBS. The following genotypes were prepped in duplicate: *w^1118^*, *pUASp-APEX2/+; nanosGAL4/+, pUASp-cona::APEX2/+; cona^f0493^/ nanosGAL4::VP16* and *pUASp-APEX2::c(2)m/ nanosGAL4::VP16.* Nuclei were isolated from ovary samples using a Singulator 100 (S2 Genomics) using low volume conditions. Nuclei samples were refrozen for future use.

To pull-down biotinylated proteins, nuclei samples were spun down at 600 x g for 10 minutes, and the solution was removed. Pelleted nuclei were ground with an electric pestle in 250 ul RIPA buffer supplemented with freshly added 5 mM Trolox and 10 mM ascorbic acid. After grinding, the solution volume was brought up to 1000 ul with the RIPA buffer solution and treated with 10 U benzonase (Millipore Sigma) at 4°C for 2 hours while rocking. Samples were spun down at max speed (20817 x g) for 15 min and the supernatant removed. 515 µL of fresh supplemented RIPA buffer was added and 15 uL was saved. The remaining 500 µL was added to 200 ul magnetic streptavidin beads (Pierce streptavidin magnetic beads 88817, Thermo Fischer Scientific) that were prepared by washing 2X in RIPA buffer. Samples were rocked overnight at 4°C. The next day, the beads were pelleted using a magnetic stand, and the solution removed and frozen. Beads were washed 2X with RIPA for 10 min at 4°C. Beads then underwent the following washes for 10 min each at 4°C: 1X 1M KCL, 1X 0.1 Na2CO3, 3X 2 M Urea in 10 mM Tris made fresh, and 2X PBS. Samples were submitted for proteomics analysis in 500 uL of fresh PBS.

### Sample preparation for mass spectrometry

Proteins bound to magnetic beads were washed twice with 500 µl of 100 mM Tris (pH 8.8). Elution was performed by incubating the beads with 60 µl of 100 mM Tris (pH 8.8) containing 2 M urea and 0.5 µg trypsin at 27 °C for 30 min with shaking. Beads were immobilized on a magnetic rack, and the supernatant was transferred to a clean microcentrifuge tube. The beads were then rinsed with 50 µl of 100 mM Tris (pH 8.8) containing 2 M urea and 1 mM tris(2-carboxyethyl)phosphine (TCEP), and this rinse was combined with the initial eluate. The combined eluates were incubated for 30 min at room temperature, after which chloroacetamide (CAM) was added to a final concentration of 10 mM. Samples were further incubated in the dark for 30 min at room temperature. Calcium chloride was then added to a final concentration of 2 mM, and digestion proceeded overnight at 37 °C. Reactions were quenched with formic acid to 5% (final concentration), centrifuged at 20,000 × g for 30 min, and the supernatants transferred to fresh tubes.

### Mass spectrometry analysis

Samples were analyzed by MudPIT, essentially as described previously (Swanson *et al*. 2009). The digested samples were loaded onto fused silica microcapillary columns containing three sequential chromatography phases: reverse phase, strong cation exchange, and reverse phase. Peptides were eluted stepwise using an Agilent 1260 Infinity quaternary HPLC pump with varying mixtures of buffer A (5% acetonitrile, 0.1% formic acid), buffer B (80% acetonitrile, 0.1% formic acid), and buffer C (5% acetonitrile, 500 mM ammonium acetate, 0.1% formic acid). Eluted peptides were introduced directly into an Orbitrap Elite mass spectrometer operated in positive ion mode, and data were collected over ten chromatography steps, each lasting two hours.

MS2 peak list files were generated from the resulting RAW files using the in-house software RawDistiller (version 1.0). Peak list files were analyzed with ProLuCID version 1.3.3, using mass tolerances of 10 ppm for precursor ions and 500 ppm for fragment ions. The sequence database used for searching contained sequences for 22,279 non-redundant *D. melanogaster* proteins downloaded from the NCBI on 2023-08-01, sequences for 426 common contaminants, 3 sequences corresponding to the recombinant APEX2 tagged proteins, and shuffled versions of all sequences for estimating false discovery rates (45,416 total sequences). Searches included differential mass shifts of 15.9949 on methionine residues (methionine oxidation) and 361.14601 on tyrosine residues (biotin-phenol modification), and a static mass shift of 57.02146 on cysteine residues (carbamidomethylation).

Following database searching, the resulting SQT files were analyzed using DTASelect (v1.9) in combination with the in-house software swallow (https://github.com/tzw-wen/kite) to control false discovery rates (FDRs), which were maintained below 5% at both the peptide and protein levels. Replicate analyses were compared to controls using Contrast (v1.9) alongside the in-house software sandmartin (https://github.com/tzw-wen/kite). The resulting Contrast.txt files were further processed with the in-house tool NSAF7x64 to generate reports containing label-free quantitation (dNSAF) values for proteins (https://github.com/tzw-wen/kite/tree/master/windowsapp/NSAF7x64). Finally, the statistical tool QPROT was applied to identify proteins significantly enriched (log2FC > 2, Z statistic > 5) in experimental samples relative to controls (Choi *et al*. 2015). Mass spectrometry data are available from the MassIVE repository (https://massive.ucsd.edu/) using the accession number MSV000098864. Table S1-3 provides the list of enriched proteins in *cona::APEX2*, *APEX2::c(2)M* and *w^1118^*samples compared to *APEX2* samples. Tables S5-6 provides additional parameters associated with the mass spectrometry experiment.

### Immunofluorescence

To examine ovaries that underwent biotin-phenol labeling by immunofluorescence, after the labeling reaction was quenched, ovaries were fixed in 4% paraformaldehyde PBT (PBS plus 0.3% Triton-X and 0.5% bovine serum albumin (BSA) (EMD Chemicals, San Diego, CA)) for 10 min. Ovaries were washed 3X with PBT for 15 minutes followed by a 4^th^ PBT for 1 hour and late-stage egg chambers were manually removed. Primary antibodies were applied overnight in PBT. The next day, ovaries were washed 3X in PBT, followed by application of secondaries in PBT for 2-4 hours. In the last 15 minutes of secondary incubation, 4’6-diamidindino-2-phenylindonle (DAPI) was added at a concentration of 1 ug/ml. After 3 washes in PBT egg chambers were mounted in ProLong Gold (Life Technologies, Grand Island NY).

The immunofluorescence of germline expressed RNAi knockdown lines was conducted as described in Hughes et al. 2025 (Hughes *et al*. 2025).

Primary antibodies used include mouse anti-C(3)G 1A8-1G2 (1:500)(Anderson *et al*. 2005), anti-γH2Av (1:5000)(Rockland Lot 46042), rat-anti-CID (1:2000-1:3000)(Hanlon *et al*. 2018), rabbit anti-Cid (1:2000)(gift of Dr. Gregory Rogers), affinity-purified rabbit anti-Corolla (1:2000) (Collins *et al*. 2014), mouse anti-Orb 4H8 and 6H4 (1:40 each)(Developmental Studies Hybridoma Bank, Iowa). Secondary antibodies used were goat Alexa Fluor anti-rabbit 488 and 555, anti-mouse 488 or 555, anti-mouse IgG1 488 or 555, anti-mouse IgG2A 647, and rat 488 (ThermoFisher). Streptavidin Alexa Fluor 488 conjugate (1:400) (S32354 Invitrogen) was added with the secondary antibodies.

### Microscope and image processing

Images were acquired with a Deltavision Elite system from GE Healthcare equipped with an Olympus IX70 inverted microscope and a high-resolution CCD camera was used. Images were deconvolved with SoftWoRx v. 7.2.1 (Applied Precision/Leica Microsystems, Inc,). SoftWoRx v. 7.2.1 or Fiji were used for image analysis. Brightness and contrast were adjusted minimally for improved publication quality.

### Scoring

For the screening of RNAi lines targeting proteins enriched in the *cona::APEX2* or *APEX2::c(2)M* expressing ovaries a minimum of 10 germaria were scored visually on the microscope for the proper assembly of full-length SC in region 2A of the germarium, maintenance of full-length SC in a single nucleus at region 3, presence of γH2Av foci in region 2A, and absence of γH2Av in the SC positive nucleus at region 3. If defects were detected in any germaria, images were acquired for future analysis. Lines had to display defects in at least 50% of scored germaria to be classified as having a defect. Lines were classified as SC assembly defect if few or no nuclei produced robust full-length SC in region 2A (this could include failure due to developmental delays). A line was classified as SC maintenance defect if full-length SC was formed in region 2A but the nucleus at region 3 displayed discontinuities in the SC tracks or more severe SC disassembly. If no γH2Av foci were present in regions 2A or 2B, a line would have been classified as having a DSB initiation defect. The presence of γH2Av foci in region 3 led to a classification of DNA repair defects. No lines caused defects only in γH2Av foci. Underdeveloped ovaries were assigned to lines in which few, if any, mid-prophase/late-stage egg chambers were present in the ovary, making the lines difficult to examine cytologically using the methods described in Immunofluorescence Materials and methods.

To score for centromere clustering, ovaries were stained with antibodies to C(3)G (SC), CID (Centromeres), and Orb (oocyte selection in later regions where the SC was disassembling in RNAi knock-down conditions). The number of CID foci that overlapped the SC or were within the nuclear region devoid of Orb staining and overlapping DAPI signal were counted for each region. For the *Cps5 RNAi* knockdown line, some germaria did not clearly have nuclei at all three early to mid-pachytene stages, and assignment of region was based on cyst location and shape.

To score for γH2Av foci number at regions 2A, 2B, and 3, ovaries were stained with antibodies to C(3)G (SC), γH2Av (DSBs), and Orb (oocyte selection). Typically, more than one region 2A nucleus was analyzed for each germarium if multiple, unobstructed nuclei were identified. If Orb had accumulated around a single nucleus in a 2B cyst, that was the nucleus scored. A nucleus was scored for γH2Av number from both cysts if more than one cyst was present in region 2B. For region 3, the Orb selected nucleus was scored for DSBs when possible. To score γH2Av foci, a nucleus was duplicated in X, Y, and Z in Fiji, and the individual Z-slices made into a montage using a Make Montage Plugin for the SC and γH2Av channels. γH2Av foci were counted as they appeared through the Z slices. Assessment of stage was based on cyst location and shape, with some *Cpsf5 RNAi* germaria not presenting a clear nucleus to score at all three stages.

### Meiotic nondisjunction and recombination assays

The frequency of meiotic nondisjunction of the *X* chromosome was measured by crossing single virgin females to *y sc cv v f car / B^s^Y* males. Nondisjunctional offspring from the mother were recovered as *B^s^* females (diplo-*X* exceptions) and *B+* males (nullo-*X* exceptions) while normal segregation results in *B+* females and *B^s^* males. The adjusted total accounts for inviable progeny classes and equals exceptional progeny X2 plus normal progeny. Nondisjunction frequency was calculated as the sum of exceptional progeny X 2 divided by adjusted total progeny. The statistical test used is described in (Zeng *et al*. 2010).

For *2^nd^* chromosome recombination analysis, single female virgin *net dpp ^d-ho^ dpy Adc^b1^ pr cn /+; nanosGAL4::VP16/+ or net dpp ^d-ho^ dpy Adc^b1^ pr cn/+; cpsf5 RNAi/nanosGAL4::VP16* were crossed to *net dpp ^d-ho^ dpy Adc^b1^ pr cn* males. Phenotypic markers were only scored in the female progeny.

### Statistics and tetrad calculations

To compare between genotypes the difference for each interval in recombination experiments Fisher’s exact test was used (https://www.graphpad.com/quickcalcs/contingency1/). The Mann-Whitney U test was used to compare genotypes for γH2Av foci number and centromere clustering (https://miniwebtool.com/mann-whitney-u-test-calculator/). The shiny app https://simrcompbio.shinyapps.io/CrossoverMLE/ was used to calculate maximum likelihood estimates (MLE) of tetrad classes (Hawley *et al*. 2025).

## Data availability

Original data underlying this manuscript can be accessed from the Stowers Original Data Repository at http://stowers.org/research/publications/libpb-2578. The *Drosophila* RNAi stocks used in this study were created by the Harvard Transgenic RNAi project (TRiP collection)(Zirin *et al*. 2020) and can be ordered from the Bloomington Stock Center. Mass spectrometry data are available from the MassIVE repository (https://massive.ucsd.edu/) using the accession number MSV000098864.

## Acknowledgments

We thank the Transgenic RNAi Project at Harvard Medical School (NIH/NIGMS RO1-GM08497) for providing transgenic RNAi fly stocks used in this study and the Bloomington Stock Center (NIH-P40D018537) for the distribution of stocks. We would like to thank Cathy Lake, Katherine K. Billmyre, and Jeremy Burton for providing insightful comments on the manuscript.

## Supplemental files

**Table S1. Proteins enriched in APEX2::C(2)M samples compared to APEX2 samples by mass spectrometry.** Enriched proteins are highlighted in yellow. Gene symbols that are in bold were identified as being enriched in both APEX2::C(2)M and Cona::APEX2 samples.

**Table S2. Proteins enriched in Cona::APEX2 samples compared to APEX2 samples by mass spectrometry.** Enriched proteins are highlighted in yellow. Gene symbols that are in bold were identified as being enriched in both APEX2::C(2)M and Cona::APEX2 samples.

**Table S3. Proteins enriched in *w^1118^* samples compared to APEX2 samples by mass spectrometry.** Enriched proteins are highlighted in yellow.

**Table S4. Results for TriP lines tested for early female meiotic phenotypes.** Descriptions are based on FlyBase and may not include all listed functions of the protein (Gramates *et al*. 2022).

**Table S5. Detailed protein list generated from all samples. Table S6. Mass spectrometry DTASelect parameters.**

